# A Combinatorial MAP Code Dictates Polarized Microtubule Transport

**DOI:** 10.1101/731604

**Authors:** Brigette Y. Monroy, Tracy C. Tan, Janah May Oclaman, Jisoo S. Han, Sergi Simo, Dan W. Nowakowski, Richard J. McKenney, Kassandra M. Ori-McKenney

## Abstract

Many eukaryotic cells distribute their intracellular components through asymmetrically regulated active transport driven by molecular motors along microtubule tracks. While intrinsic and extrinsic regulation of motor activity exists, what governs the overall distribution of activated motor-cargo complexes within cells remains unclear. Here, we utilize in vitro reconstitution of purified motor proteins and non-enzymatic microtubule-associated proteins (MAPs) to demonstrate that these MAPs exhibit distinct influences on the motility of the three main classes of transport motors: kinesin-1, kinesin-3, and cytoplasmic dynein. Further, we dissect how combinations of MAPs affect motors, and reveal how transient interactions between MAPs and motors may promote these effects. From these data, we propose a general “MAP code” that has the capacity to strongly bias directed movement along microtubules and helps elucidate the intricate intracellular sorting observed in highly polarized cells such as neurons.

## INTRODUCTION

Within cells, nothing works in isolation. Therefore, in order to dissect the complexity of intracellular processes, it is essential to study the behaviors of molecules both individually and collectively. One such intricate process is the polarized active transport along microtubules that is required within neurons for the establishment and maintenance of distinct dendritic and axonal compartments ^1,2^. This efficient transport system is driven by kinesin motors and cytoplasmic dynein, which travel towards the microtubule plus and minus ends, respectively ^2–4^. Of the large kinesin family, the kinesin-1, -2, and -3 classes are thought to act as the predominant long-distance transport motors, while other kinesins serve more specialized cellular roles ^5,6^. In neurons, kinesin-3 and dynein drive various cargoes within both axons and dendrites, while kinesin-1 carries cargoes into the axon, but is largely excluded from the dendrites ^6–10^. Post-translational modifications of tubulin have been proposed to act as a “tubulin-code” that can be read out by activated motor proteins to direct their movement to specific cellular compartments ^9^. However, other than the effect of tyrosination on dynein landing rate ^11^, the reported biophysical effects of certain tubulin modifications, such as acetylation, on kinesin motor movement are relatively modest ^12^, raising questions about how such effects could directly result in guiding transport in vivo. A large variety of other proteins bind to microtubules, and as such, transport motors must encounter a number of non-enzymatic microtubule-associated proteins (MAPs) that decorate the microtubule cytoskeleton ^13^. Disruption of this bidirectional transport system due to mutations in motor complexes or MAPs leads to a wide range of neurodevelopmental and neurodegenerative disorders ^13–17^, highlighting the interplay between these classes of proteins.

Since the identification of “structural” MAPs that co-purified with polymerized brain tubulin ^18^, such as MAP1, MAP2, tau, MAP7, and doublecortin (DCX), MAPs have been described as stabilizers, nucleation-promoting factors, and bundlers of microtubules ^19–24^. However, recent work suggests that these MAPs may also function to direct motor transport ^6,25–27^. Perhaps the most well-studied MAP with regards to its effects on motors is the Alzheimer’s disease-associated MAP, tau, which was originally thought to be axon-specific, but can also be observed in mature dendrites (Figure S1) ^27,28^. Tau inhibits kinesin-1 and kinesin-3 to varying degrees ^25,26,29–31^, but does not strongly impede processive dynein motility ^27^. These differential effects are due to a steric clash between tau and the relatively large kinesin motor domain, which does not exist for the smaller dynein microtubule-binding domain ^27,32,33^. MAP2 is localized to dendrites and the axon initial segment and has been shown to inhibit kinesin-1 in vivo ^34^. Based on the similarities in their microtubule binding domains and binding sites on the lattice ^32,35^, it is likely that MAP2 affects kinesins and dynein akin to tau. MAP7 is important for a range of kinesin-1 functions in vivo ^36–40^, and has been shown in vitro to directly bind and recruit kinesin-1 to the microtubule lattice ^26,41,42^. Kinesin-1 is most likely able to navigate the tau-rich axon in part due to the presence of MAP7, which displaces tau from the microtubule ^26^. Interestingly, MAP7 inhibits kinesin-3, but does not substantially affect dynein motility ^26^. DCX and its paralogue, doublecortin-like kinase-1 (DCLK1) robustly stimulate microtubule polymerization ^23,43,44^, but are restricted to distal dendrites and axonal growth cones ^6,45,46^, indicating they may have specific roles commensurate with their localization patterns. Both MAPs have been reported to interact with the kinesin-3 motor domain and promote kinesin-3 cargo transport within dendrites ^6,46^; however, the molecular mechanism underlying this relationship is unclear. Another interesting MAP is the understudied MAP9/ASAP, which plays a role in organizing the mitotic spindle in cultured cells ^47,48^, and is associated with cell degeneration and cancer ^49,50^. Although MAP9 is highly expressed in the vertebrate nervous system throughout development ^51^, its molecular function is unknown. Why there are distinct, yet overlapping localization patterns for these MAPs within neurons and how these MAPs may contribute, individually and collectively, to sorting motors into specified compartments remain outstanding questions.

Here, we present a comprehensive analysis of the effect of six MAPs on three classes of transport motors in an effort to elucidate a general “MAP code” that could underlie polarized transport. We find that tau and MAP2 act as general inhibitors of kinesin-1 and kinesin-3, preventing these motors from accessing the lattice, while three MAPs that localize within dendrites, DCX, DCLK1, and MAP9 differentially gate access to the microtubule by inhibiting kinesin-1, but not kinesin-3, providing a molecular system by which these motors are spatially regulated in neurons. We dissect the mechanism by which kinesin-3 is able to progress through these MAPs, highlighting a key role for MAP9 in specifically facilitating kinesin-3 translocation. Furthermore, MAP9 is the only neuronal MAP examined thus far that substantially inhibits the processive cytoplasmic dynein complex. Overall, our study provides general mechanistic principles for how MAPs help to orchestrate the distribution of specific motors within the crowded intracellular environment by gating access to the microtubule lattice.

## RESULTS

### Compartmentally distinct MAPs differentially affect kinesin-1 and kinesin-3

In order to understand how motors are differentially directed into dendritic or axonal compartments, we first wanted to determine the localization patterns of six MAPs within neurons at the same developmental time point. We performed immunocytochemistry on DIV4 primary mouse hippocampal cultures with antibodies against tau, MAP2, MAP7, DCX, DCLK1, and MAP9 (Figure S1). We found that three of these MAPs, tau, MAP7, and MAP9, localized throughout both dendrites and axons, while the other three, MAP2, DCLK1 and DCX, were predominantly restricted to dendrites, consistent with prior localization studies (Figure S1) ^6,26–28,45,46^.

Because of the spatial distributions of kinesin family proteins, we set out to investigate how compartmentally distinct MAPs may affect kinesin-1 and kinesin-3 motility along microtubules in an effort to understand how MAPs, in general, could contribute to polarized motor transport in vivo. Towards this goal, we utilized a molecular reconstitution system of purified proteins to probe for direct effects of MAPs on motor motility. Using multi-color total internal reflection fluorescence microscopy (TIRF-M), we imaged the progression of purified, fluorescently labeled truncated kinesin motors, K560 (kinesin-1, KIF5B_1-560_) and KIF1a_1-393_ (kinesin-3), in the absence or presence of six fluorescently labeled full-length MAPs (Figures 1-2 and S2A-B). Strikingly, we observed that other than MAP7, which increased kinesin-1 landing rate 25-fold ^26^, all other MAPs significantly decreased the landing rate of kinesin-1 on the microtubule lattice, with the greatest effect being a 15-fold reduction by MAP2 (Figure 1A-B). It was especially surprising that DCX and DCLK1 both inhibited kinesin-1, considering they do not share overlapping binding sites with the kinesin-1 motor domain ^43^, suggesting this effect may be due to steric interference away from the surface of the MT. With the exception of MAP7, all other MAPs present in the dendrites blocked kinesin-1 from landing on the microtubule, suggesting that the mere presence of MAP7 may not be sufficient to facilitate kinesin-1 transport in all cellular compartments. To test this idea, we asked if MAP7 could facilitate kinesin-1 motility in the presence of an inhibitory MAP (iMAP). Both DCX and MAP9 bind simultaneously with MAP7 on microtubules, suggesting their binding sites do not overlap (Figure 1C). In the presence of either of these iMAPs and MAP7, we found that although kinesin-1 was recruited to the microtubule by MAP7, its movement along the lattice was still largely inhibited (Figure 1B-D). Thus, diverse iMAPs have dominant effects on kinesin-1 movement even after the motor has been recruited to the microtubule surface by MAP7.

**Figure 1.**
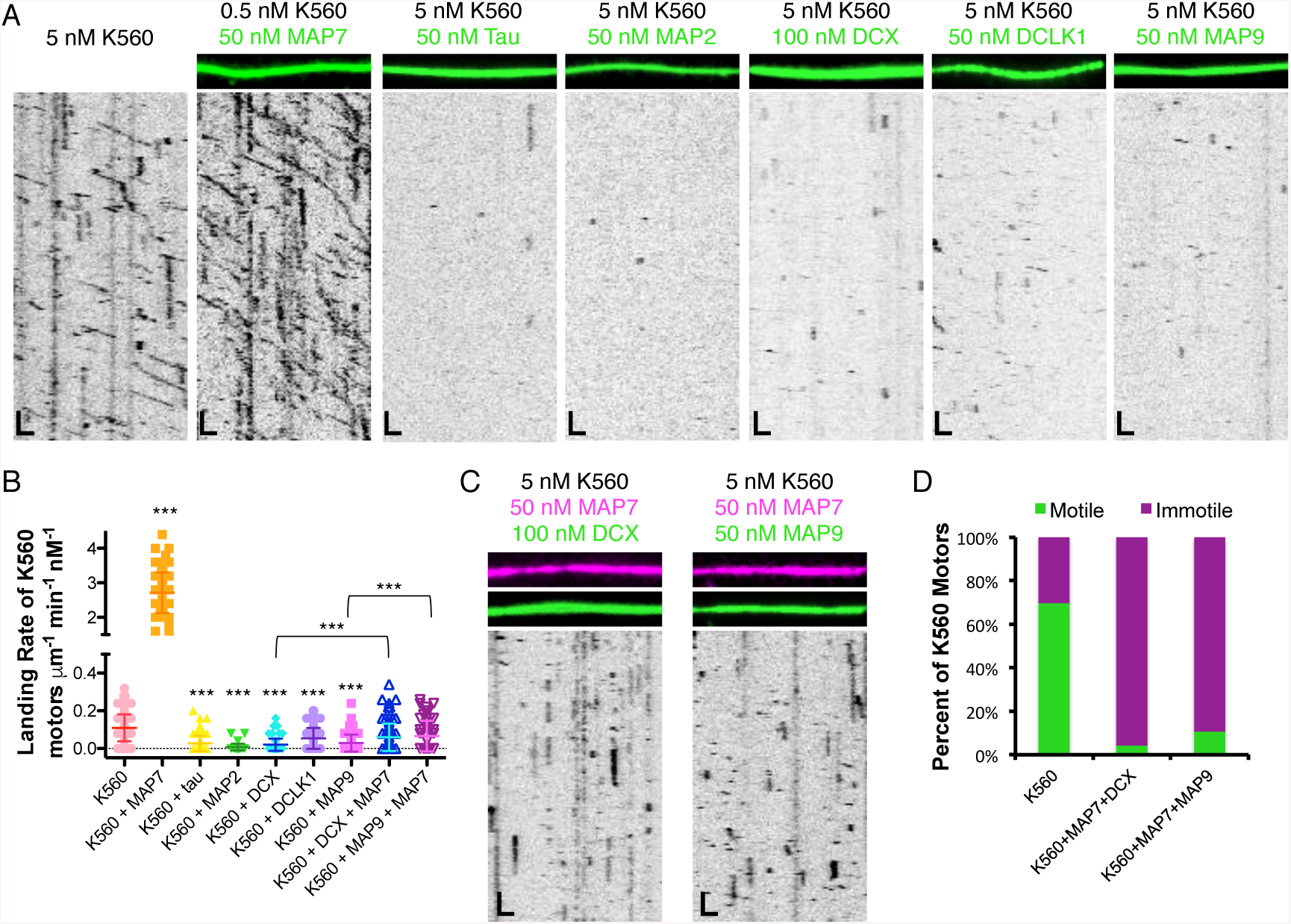
Kinesin-1 is inhibited from landing and progressing on the microtubule lattice by tau, MAP2, DCX, DCLK1, and MAP9. **(A)** TIRF-M images and kymographs of K560-mScarlet (kinesin-1) at indicated concentrations + 1 mM ATP in the absence and presence of 50 nM sfGFP-MAP7, 50 nM sfGFP-tau, 50 nM sfGFP-MAP2, 100 nM DCX-sfGFP, 50 nM sfGFP-DCLK1, or 50 nM sfGFP-MAP9 (green). Images are 10.3 μm wide. Scale bars: 1 μm (*x*) and 7 sec (*y).* **(B)** Quantification of the landing rates of K560-mScarlet + 1 mM ATP in the absence and presence of each MAP or MAP combination (means ± s.d. in motors μm^-1^ min^-1^ nM^-1^ are: 0.11 ± 0.07 for K560 alone (n = 134 kymographs from three independent trials), 2.72 ± 0.59 for K560 + MAP7 (n = 83 kymographs from two independent trials), 0.03 ± 0.04 for K560 + tau (n = 100 kymographs from two independent trials), 0.01 ± 0.02 for K560 + MAP2 (n = 93 kymographs from two independent trials), 0.02 ± 0.03 for K560 + DCX (n = 94 kymographs from two independent trials), 0.05 ± 0.05 for K560 + DCLK1 (n = 92 kymographs from two independent trials), 0.03 ± 0.05 for K560 + MAP9 (n = 96 kymographs from two independent trials), 0.06 ± 0.07 for K560 + MAP7 + DCX (n = 114 kymographs from two independent trials), and 0.07 ± 0.08 for K560 + MAP7 + MAP9 (n = 70 kymographs from two independent trials). All datapoints are plotted with lines indicating means ± s.d. *P* < 0.0001 (***) using a student’s t-test for K560 alone *vs.* K560 + each MAP individually, K560 + DCX *vs.* K560 + DCX + MAP7, and K560 + MAP9 *vs.* K560 + MAP9 + MAP7. **(C)** TIRF-M images and kymographs of 5 nM K560-mScarlet + 1 mM ATP in the presence of 50 nM BFP-MAP7 (pink) with 100 nM DCX-sfGFP (green), or 50 nM BFP-MAP7 (pink) with 50 nM sfGFP-MAP9 (green). Images are 10.3 μm wide. Scale bars: 1 μm (*x*) and 7 sec (*y).* **(D)** Quantification of the percentage of K560 motors that are motile *vs.* immotile in the absence (n = 169 molecules quantified from three independent trials) or presence of 50 nM MAP7 with 100 nM DCX (n = 160 molecules quantified from two independent trials), or 50 nM MAP7 with 50 nM MAP9 (n = 111 molecules quantified from two independent trials).

**Figure 2.**
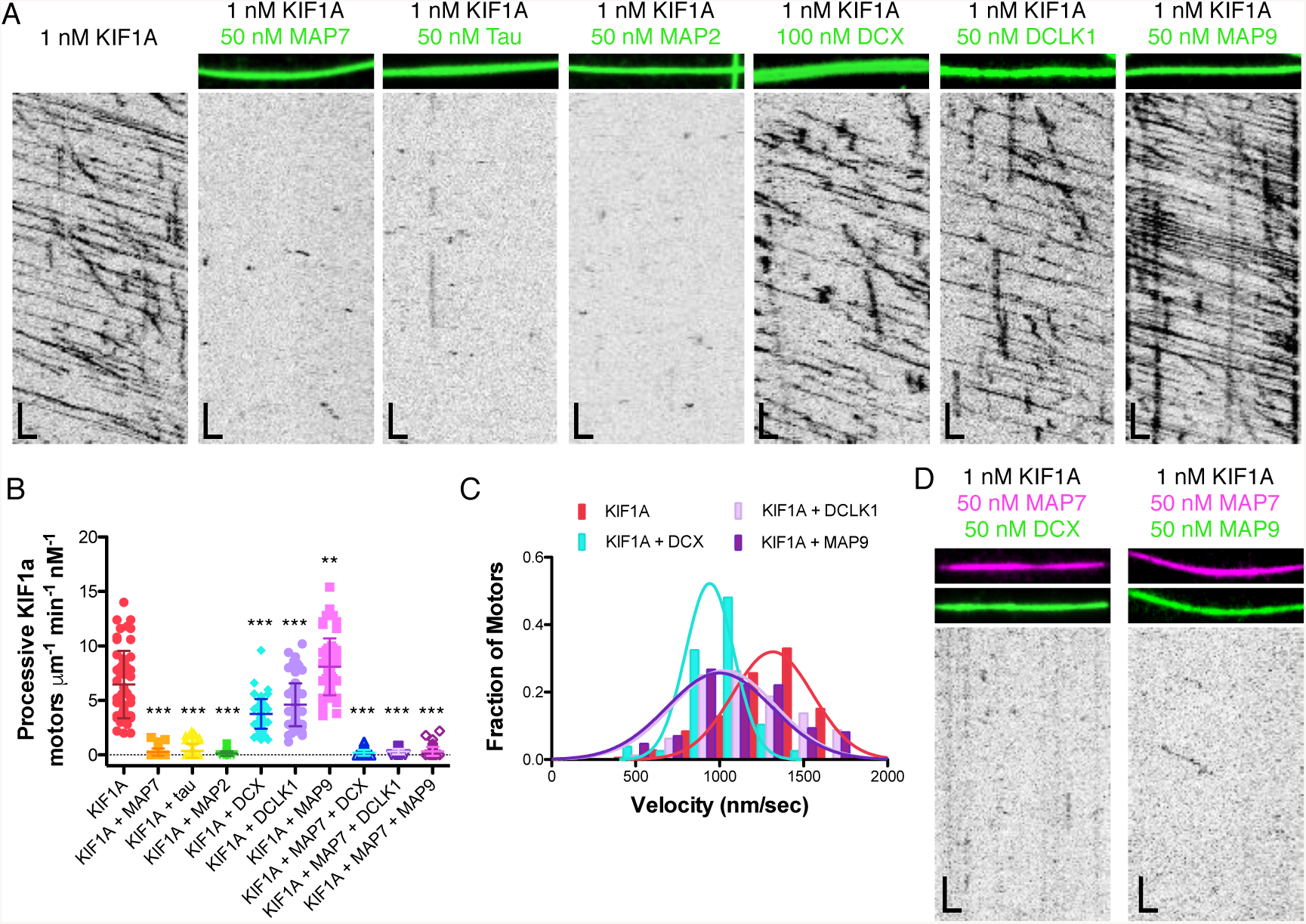
Kinesin-3 access to the microtubule lattice is differentially gated by MAP7, tau, MAP2, DCX, DCLK1, and MAP9. **(A)** TIRF-M images and kymographs of 1 nM KIF1A-mScarlet (kinesin-3) + 1 mM ATP in the absence and presence of 50 nM sfGFP-MAP7, 50 nM sfGFP-tau, 50 nM sfGFP-MAP2, 100 nM DCX-sfGFP, 50 nM sfGFP-DCLK1, or 50 nM sfGFP-MAP9 (green). Images are 10.3 μm wide. Scale bars: 1 μm (*x*) and 5 sec (*y).* **(B)** Quantification of the number of processive KIF1A-mScarlet motors + 1 mM ATP in the absence and presence of each MAP or MAP combination (means ± s.d. in motors μm^-1^ min^-1^ nM^-1^ are: 6.46 ± 3.09 for KIF1A alone (n = 54 kymographs from three independent trials), 0.26 ± 0.33 for KIF1A + MAP7 (n = 75 kymographs from three independent trials), 0.36 ± 0.64 for KIF1A + tau (n = 71 kymographs from three independent trials), 0.13 ± 0.21 for KIF1A + MAP2 (n = 69 kymographs from three independent trials), 3.77 ± 1.36 for KIF1A + DCX (n = 65 kymographs from three independent trials), 4.61 ± 1.97 for KIF1A + DCLK1 (n = 83 kymographs from three independent trials), 8.10 ± 2.61 for KIF1A + MAP9 (n = 55 kymographs from three independent trials), 0.19 ± 0.26 for KIF1A + MAP7 + DCX (n = 66 kymographs from two independent trials), 0.30 ± 0.25 for KIF1A + MAP7 + DCLK1 (n = 46 kymographs from two independent trials),and 0.33 ± 0.40 for KIF1A + MAP7 + MAP9 (n = 76 kymographs from two independent trials). All datapoints are plotted with lines indicating means ± s.d. *P* < 0.0001 (***) using a student’s t-test for KIF1A alone *vs.* KIF1A + MAP7, tau, MAP2, DCX, and DCLK1 and for each MAP combination. *P* = 0.0034 (**) for KIF1A alone *vs.* KIF1A + MAP9. **(C)** Velocity histograms of KIF1A + 1 mM ATP in the absence and presence of DCX, DCLK1, and MAP9 with Gaussian fits. Means ± s.d. are 1252.5 ± 279.7, 931.0 ± 177.0, 1070.6 ± 280.6, and 1023.8 ± 293.6 nm/sec for KIF1A alone, KIF1A + DCX, KIF1A + DCLK1, and KIF1A + MAP9, respectively. *P* < 0.0001 using a student’s t-test for KIF1A alone *vs.* KIF1A with each MAP; n = 179, 77, 80 and 86 KIF1A motors for KIF1A alone, KIF1A + DCX, KIF1A + DCLK1, and KIF1A + MAP9, respectively from two independent trials each. **(D)** TIRF-M images and kymographs of 1 nM KIF1A-mScarlet + 1 mM ATP in the presence of 50 nM BFP-MAP7 (pink) with 50 nM DCX-sfGFP (green), or 50 nM BFP-MAP7 (pink) with 50 nM sfGFP-MAP9 (green). Images are 10.3 μm wide. Scale bars: 1 μm (*x*) and 5 sec (*y).*

Considering the similarity in motor domains and binding footprints on the microtubule lattice between kinesin-1 and kinesin-3 ^52,53^, we were curious if we would observe the same global inhibition of kinesin-3 by these six MAPs. Indeed, we observed that MAP7, tau, and MAP2 largely inhibited kinesin-3 from accessing the microtubule. Strikingly, saturating amounts of DCX, DCLK1, and MAP9 were permissive for kinesin-3 motility (Figure 2A-B). Prior studies have reported that DCX and DCLK1 interact with the motor domain of kinesin-3 and are important for kinesin-3 transport of cargo within dendrites in vivo ^6,46^. However, we did not observe an increase in kinesin-3 landing rate or motor velocity in the presence of DCX or DCLK1 (Figure 2A-C), indicating that these MAPs do not directly recruit kinesin-3 or allosterically enhance its motor activity. Interestingly, MAP9 was the only MAP to significantly increase the number of processive kinesin-3 motors on the lattice (Figure 2A-B). We next wanted to test if DCX, DCLK1, and MAP9 could recruit kinesin-3 or promote kinesin-3 motility in the presence of an iMAP. On microtubules co-decorated with saturating amounts of MAP7 and either DCX, DCLK1, or MAP9, we still observed a significant inhibition of kinesin-3 landing events (Figure 2B,D). Thus, unlike MAP7, which can recruit kinesin-1 in the presence of an iMAP, we do not observe a similar effect for DCX, DCLK1, or MAP9 on kinesin-3, suggesting these MAPs do not stably interact with the kinesin-3 motor domain. However, our data indicate that MAP9, which is present in both dendrites and axons (Figure S1), is the only MAP that facilitates kinesin-3 motility along the microtubule.

### MAP9 enables kinesin-3 progression on the lattice due to a transient charged interaction

It is striking that three MAPs present in dendrites, DCX, DCLK1, and MAP9, impede kinesin-1, but not kinesin-3. We therefore wanted to investigate the differential effects of these three MAPs on kinesin-1 and kinesin-3. At sub-saturating concentrations of DCX (5 nM), where we observe cooperative clusters of DCX molecules on the microtubule lattice ^23^, substantially more kinesin-3 motors are able to enter and pass these clusters than kinesin-1 motors, the majority of which detach upon encountering DCX assemblies (Figure S3A-B). Similar to kinesin-3, the processive dynein-dynactin-BicD (DDB) complex, which also transports cargo in the dendrites, largely moved through cooperative DCX patches unimpeded (Fig S3A-B). We, and others, have previously shown that MAP7 directly interacts with kinesin-1 to recruit kinesin-1 to the microtubule ^26,41^. Similarly, prior studies have reported a direct interaction between kinesin-3 and both DCX ^46^ and DCLK1 ^6^. We therefore asked whether a motor must directly interact with a MAP in order to progress through a MAP-decorated lattice. Solution-based pull-down assays using purified proteins revealed that neither DCLK1 nor MAP9 stably interacted with our kinesin-3 construct (KIF1A_1-393_) (Figure 3A). We cannot rule out a possible interaction between these MAPs and the kinesin-3 tail domain, which is missing in our construct; however, previous data suggested that the kinesin-3 motor domain directly interacts with these MAPs ^6,46^. Although our result is in contrast to these studies, it is consistent with our observations that DCX, DCLK1, and MAP9 are unable to recruit kinesin-3 in the presence of an iMAP, in contrast to the ability of MAP7 to recruit kinesin-1 in a similar assay (Figure 1B-C, 2B,D).

**Figure 3.**
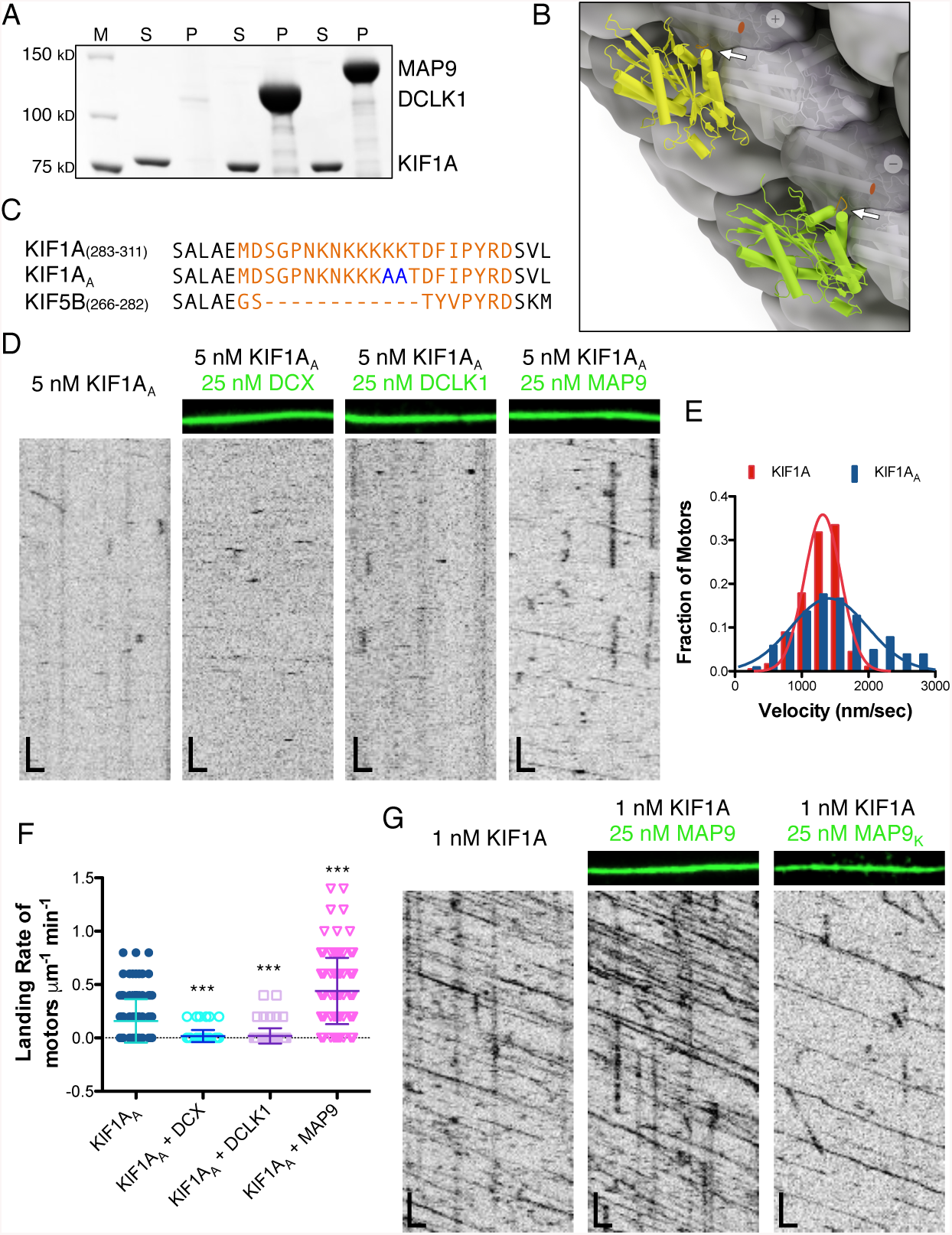
Mechanistic dissection of kinesin-3 progression through MAP-decorated microtubule lattices. **(A)** Coomassie Blue-stained SDS-PAGE gel of FLAG pull-downs with purified recombinant proteins. Either full-length sfGFP-MAP9-FLAG or sfGFP-DCLK1-FLAG were used for pull-down assays with KIF1A-mScarlet (n = three independent trials per assay). M = marker, S = supernatant, and P = pellet. **(B)** Visualization of kinesin-3 (KIF1A, PDB: 1i5s; yellow) and kinesin-1 (KIF5B, PDB: 4hna; green) motor domains on the microtubule. Arrows point to loop 12 in orange (KIF5B residues 271-279 and KIF1A residues 288-308; 14 residues of KIF1A, 290-303, are not structurally resolved). Red dots mark C-termini on beta-tubulin, where disordered C-terminal tails are not structurally resolved. **(C)** Sequence alignment comparing loop 12 of kinesin-3 (KIF1A) and kinesin-1 (KIF5B). Sequence of KIF1A_AA_ indicates the two lysines that were mutated to alanines (blue) for the studies in **(D). (D)** TIRF-M images and kymographs of 5 nM KIF1A_AA_-mScarlet (kinesin-3) + 1 mM ATP in the absence and presence of 50 nM DCX-sfGFP, 50 nM sfGFP-DCLK1, or 50 nM sfGFP-MAP9 (green). Images are 9.1 μm wide. Scale bars: 1 μm (*x*) and 5 sec (*y).* **(E)** Velocity histograms of KIF1A and KIF1A_AA_ (in the presence of MAP9) + 1 mM ATP with Gaussian fits. KIF1A data are reproduced from Figure 2C for comparison. There were too few KIF1A_AA_ motors alone to analyze, thus we quantified motor velocity in the presence of MAP9 due to the increased motor density on the microtubule. Mean ± s.d velocity for KIF1A_AA_ + MAP9 was 1503.7 ± 642.0 nm/sec (n = 102 KIF1A motors from three independent trials). *P* < 0.0001 using a student’s t-test for KIF1A *vs.* KIF1A_AA_. **(F)** Quantification of the landing rates of 5 nM KIF1A_AA_-mScarlet + 1 mM ATP in the absence and presence of DCX, DCLK1, and MAP9 (means ± s.d. in motors μm^-1^ min^-1^ are: 0.16 ± 0.20 for KIF1A_AA_ alone (n = 143 kymographs from three independent trials), 0.02 ± 0.06 for KIF1A_AA_ + DCX (n = 92 kymographs from two independent trials), 0.02 ± 0.07 for KIF1A_AA_ + DCLK1 (n = 94 kymographs from two independent trials), 0.44 ± 0.31 for KIF1A_AA_ + MAP9 (n = 135 kymographs from three independent trials). All datapoints are plotted with lines indicating means ± s.d. *P* < 0.0001 (***) using a student’s t-test for KIF1A_AA_ alone *vs.* KIF1A_AA_ + each MAP. **(G)** TIRF-M images and kymographs of 1 nM KIF1A-mScarlet + 1 mM ATP in the absence and presence of 25 nM sfGFP-MAP9 or 25 nM sfGFP-MAP9_EEE>KKK_ (green). Images are 10.3 μm wide. Scale bars: 1 μm (*x*) and 5 sec (*y)*. Mean ± s.d landing rate for KIF1A + MAP9_EEE>KKK_ was 1.22 ± 0.68 μm^-1^ min^-1^ nM^-1^ (n = 94 kymographs from two independent trials). *P* < 0.0001 using a student’s t-test for KIF1A *vs.* KIF1A + MAP9_EEE>KKK_.

A primary difference between the kinesin-1 and kinesin-3 constructs used in this study is the presence of six lysines, known as the K-loop, within loop 12 of the kinesin-3 motor domain (Figure 3B-C). We mutated two of these lysines to alanines (kinesin-3_A_) and observed the effects on motility in the absence and presence of DCX, DCLK1, and MAP9 (Figure 3C-F). Similar to prior studies of this region ^54,55^, the kinesin-3_A_ motor exhibited a significant reduction in landing rate without a decrease in velocity (Figure 3D-F). The presence of DCX or DCLK1 further decreased the landing rate of kinesin-3_A_, but the presence of MAP9 on the microtubule actually increased the landing rate of kinesin-3_A_ 3-fold (Figure 3D-F), revealing that MAP9 may directly interact with kinesin-3 on the microtubule surface, despite our inability to detect a stable interaction between the two molecules in solution. Mutating all six lysines to alanines abolished the ability of MAP9 to facilitate kinesin-3_A-loop_ landing and progression on the microtubule (Figure S4A-B), indicating that the remaining four lysines in the K-loop of kinesin-3_A_ may be sufficient to interact with MAP9. From these data, we suggest that MAP9 could transiently contact the positively charged K-loop as kinesin-3 steps along the lattice.

In order to examine a potential interaction between MAP9 and kinesin-3 on the microtubule, we examined the MAP9 sequence for a conserved acidic region that could transiently interact with the K-loop of kinesin-3 (Figure S4C-E). Mutating a conserved stretch of three glutamic acids (aa 502-504) to lysines (Figure S4D-E) strongly perturbed the ability of kinesin-3 motors to bind and move along microtubules saturated with MAP9_K_ (Figure 3G and S4F). These mutations do not increase the microtubule binding affinity of MAP9_K_ (Figure S4F), indicating that the inhibition of kinesin-3 is not due to a tighter association of MAP9_K_ with the microtubule. Collectively, these data demonstrate a mechanism by which MAP9 enables kinesin-3 to land and progress on a microtubule.

If the K-loop of kinesin-3 is the defining feature that facilitates its movement on a microtubule crowded with MAPs that inhibit kinesin-1, then the transposition of this region into kinesin-1 should confer resistance to the strong effects of its iMAPs. In support of this idea, when the kinesin-3 K-loop was inserted into loop 12 of kinesin-1 ^52,53^, the motor was able to transport cargoes into dendrites from which it is normally excluded ^56^. Based on prior studies, we engineered a chimeric kinesin-1 with the K-loop of kinesin-3 inserted into loop 12 of kinesin-1 (Figure 4A), and found that the kinesin-1 chimera (K560_K_) exhibited a 40-fold increase in landing rate compared to the wild type motor, with a small increase in velocity (Figure 4B-D). In contrast to wild-type K560, K560_K_ was able to land and translocate on a MAP9-decorated microtubule, but was impeded by the presence of MAP7, which normally inhibits kinesin-3 (Figure 4B-C). This result suggests that the K-loop of kinesin-3 is the structural element responsible for its inhibition by MAP7, because even though K560_K_ includes the MAP7 interaction region, it is still significantly inhibited from binding and moving along a lattice saturated with MAP7.

**Figure 4.**
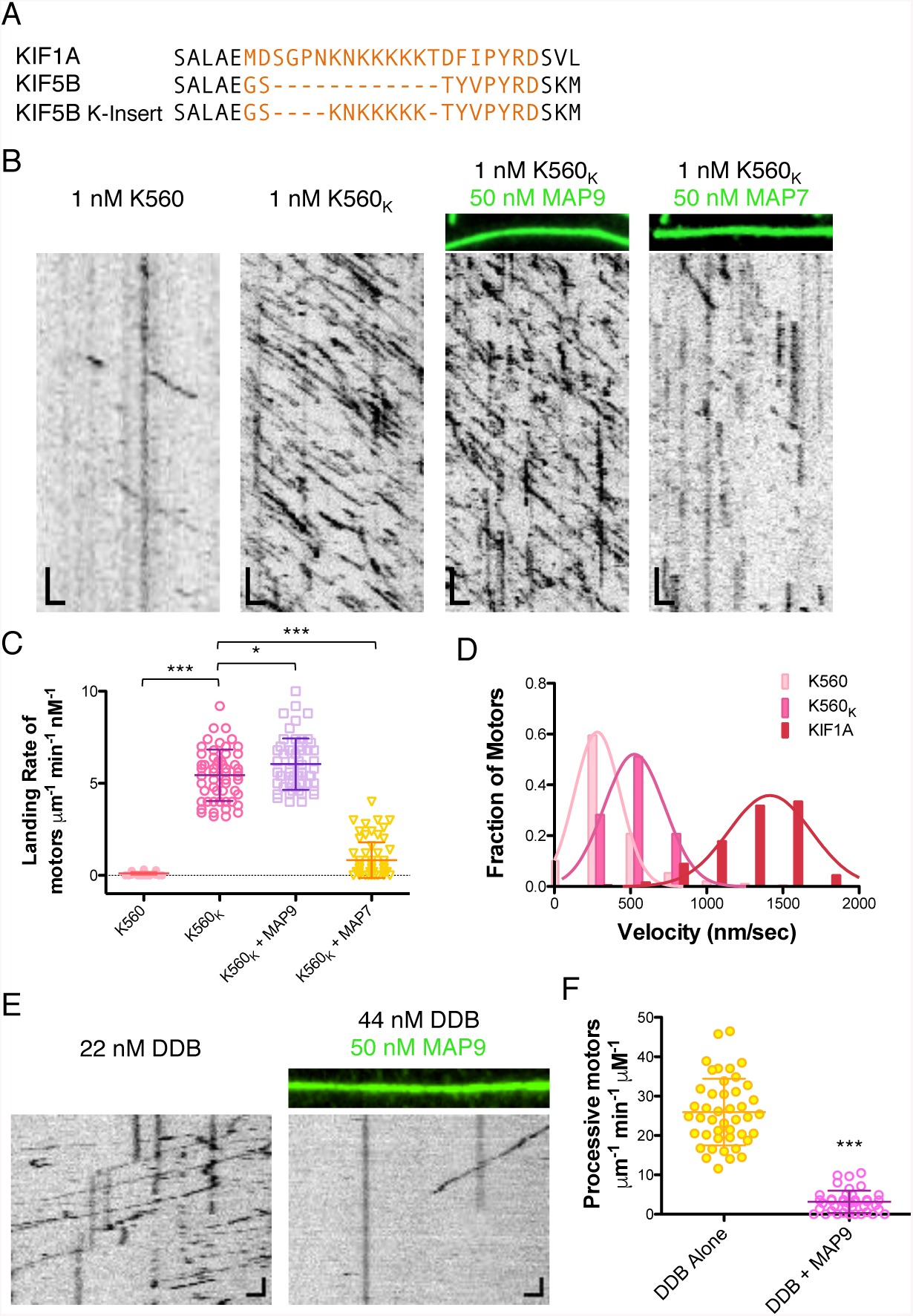
Addition of the kinesin-3 K-loop into loop 12 of kinesin-1 enhances on-rate and enables motility through a lattice saturated with MAP9. **(A)** Sequence alignment comparing loop 12 of kinesin-3 (KIF1A) and kinesin-1 (KIF5B) and the kinesin-1 chimera construct with the K-loop residues inserted into loop 12 of KIF5B for the studies in **(B). (B)** TIRF-M images and kymographs of 1 nM K560-mScarlet compared with 1 nM K560-K (kinesin-1 chimera with the K-loop insertion from kinesin-3) + 1 mM ATP in the absence and presence of 50 nM sfGFP-MAP9 or 50 nM sfGFP-MAP7 (green). Images are 10.3 μm wide. Scale bars: 1 μm (*x*) and 5 sec (*y).* **(C)** Quantification of the landing rates of K560-mScarlet compared with K560-K-mScarlet + 1 mM ATP in the absence and presence of MAP9 or MAP7. K560 data are reproduced from Figure 2B for comparison. Means ± s.d. in motors μm^-1^ min^-1^ nM^-1^ are: 5.44 ± 1.39 for K560-K alone (n = 55 kymographs from two independent trials), 6.04 ± 1.39 for K560-K + MAP9 (n = 56 kymographs from two independent trials), and 0.83 ± 0.98 for K560-K + MAP7 (n = 63 kymographs from two independent trials). All datapoints are plotted with lines indicating means ± s.d. *P* < 0.0001 (***) using a student’s t-test for K560 *vs.* K560-K, and K560-K alone *vs.* K560-K + MAP7. *P* = 0.025 (*) for K560-K *vs.* K560-K + MAP9. **(D)** Velocity histograms of K560, K560-K, and KIF1A + 1 mM ATP with Gaussian fits. KIF1A data are reproduced from Figure 3C for comparison. Mean ± s.d velocities for K560 and K560-K are 358.2 ± 295.7 nm/sec (n = 188 motors from three independent trials) and 489.0 ± 164.5 nm/sec (n = 241 motors from two independent trials), respectively. *P* < 0.0001 using a student’s t-test for K560 *vs.* K560-K. **(E)** TIRF-M images and kymographs of dynein-dynactin-BicD2 (DDB)-TMR at indicated concentrations + 1 mM ATP in the absence and presence of 50 nM sfGFP-MAP9. Images are 14.3 μm wide. Scale bars: 1 μm (*x*) and 5 sec (*y).* **(F)** Quantification of the number of processive DDB complexes + 1 mM ATP in the absence and presence of MAP9. Means ± s.d. in motors μm^-1^ min^-1^ μM^-1^ are: 26.0 ± 8.5 for DDB alone (n = 44 kymographs from two independent trials) and 3.2 ± 2.8 for DDB + MAP9 (n = 41 kymographs from two independent trials). *P <* 0.0001 (***) using a student’s t-test for DDB *vs.* DDB + MAP9.

Next, we wanted to examine the effect of MAP9 on dynein, because we observed that DCX did not significantly impair DDB translocation along the microtubule (Figure S3), and in prior studies, we have found that tau and MAP7 also do not dramatically impede dynein motility ^26,27^. In contrast to these other MAPs, MAP9 significantly inhibited processive DDB from accessing the lattice as evidenced by the 4-fold reduction in the number of processive motors on the microtubule in the presence of MAP9 (Figure 4E-F). Overall, these data reveal that MAP9 acts as a general inhibitor for kinesin-1 and dynein, but enables kinesin-3 to land and progress on the microtubule due to a transient interaction with the positively charged K-loop.

### MAP9 competes for binding on the microtubule surface with DCX and DCLK1

Very little is known about the mechanism of microtubule binding by MAP9, but we found it intriguing that, similar to DCX and DCLK1, it inhibited kinesin-1, but allowed for kinesin-3 movement. However, unlike DCX, MAP9 inhibits the association of processive dynein with the microtubule. We were therefore curious about the binding site of MAP9 on the lattice, and how its presence affected the binding of other MAPs. First, we analyzed whether MAP9 and DCX or DCLK1 could bind simultaneously to individual microtubules. We mixed equimolar concentrations of MAP9 with DCX or DCLK1 and observed that MAP9 was strongly excluded from sites of DCX or DCLK1 enrichment, and vice versa (Figure 5A). Interestingly, this anti-correlation is similar to what we observed within distal dendrites for MAP9 and DCLK1 (Figure S5). Conversely, at equimolar concentrations, MAP9 simultaneously bound microtubules coated in tau or MAP7 (Figure 5B). Our prior study on MAP7 and tau led us to hypothesize that MAP7 binds along the protofilament “ridge” of the microtubule similar to tau ^26^. DCX and DCLK1 bind the vertex of four tubulin heterodimers, occupying the “valley” between two adjacent protofilaments ^43,57,58^. Taken together, these data suggest that MAP9 also binds within the inter-protofilament valley, similar to DCX and DCLK1 (Figure 5C), but could potentially make contacts with the ridge of the protofilament to obstruct the binding domains of kinesin-1 and dynein. In addition, it is noteworthy that MAP9 can bind the microtubule simultaneously with MAP7 and tau, all three of which are present in the axon and play distinct roles in allowing kinesin-1, kinesin-3, and dynein to access the lattice.

**Figure 5:**
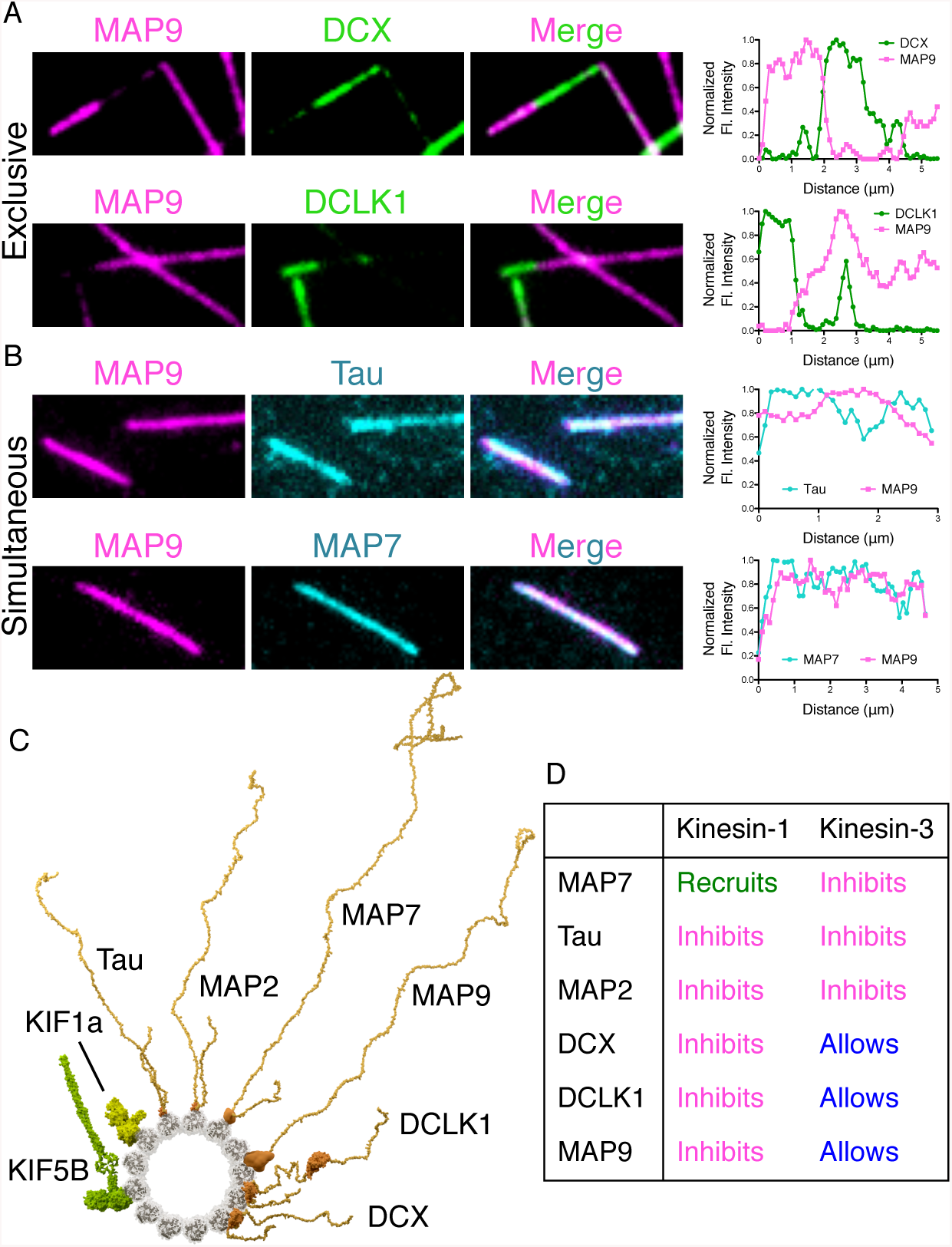
MAP9 competes for binding on the microtubule with DCX and DCLK1, but not with tau and MAP7. **(A)** TIRF-M images of 50 nM sfGFP-MAP9 and 50 nM mTagBFP-DCX or 50 nM sfGFP-MAP9 and 50 nM TagRFP-DCLK1 show these MAPs exclude each other into homotypic patches on microtubules. Right: corresponding line scans depicting fluorescence intensity of MAP9 and DCX or MAP9 and DCLK1 on a co-decorated microtubule. **(B)** TIRF-M images of 50 nM sfGFP-MAP9 and 50 nM mScarlet-tau or 50 nM sfGFP-MAP9 and 50 nM mTagBFP-MAP7 show these MAPs bind to microtubules simultaneously. Right: corresponding line scans depicting fluorescence intensity of MAP9 and tau or MAP9 and MAP7 on a co-decorated microtubule. **(C)** Model of MAPs and kinesin motor constructs bound to microtubule. End-on view of tubulin dimers is shown from the minus (-) end. KIF5B, 1-560 construct: KIF5B motor domains (4hna), and based on alignment to dimeric kinesin (3kin), coiled-coil (1d7m) as in Zhang et al. (2012). KIF1A, 1-393 construct: motor domains (1ia0), and processive dimer based on conventional kinesin (3kin) as in Huo et al. (2012). Tau: 2N4R (6cvn). MAP2: microtubule binding region based on tau (6cvn). MAP7 and MAP9: plausible microtubule-binding regions via *de novo* and homology modeling (Buchan et al., 2010). DCLK1: microtubule-binding N-DC1 domain (1mg4) and kinase domain (5jzj). DCX: microtubule-bound N-DC1 domain (2xrp). C-terminal DC2 domains are not visible for DCX and DCLK1. MAPs are shown as full-length pseudo-models, with projection domains (and domain linkers) modeled as unfolded, to visualize their length organizational differences, and to convey intrinsic disorder predicted for those regions. **(D)** Summary table of results. MAPs have distinct effects on different motors.

## DISCUSSION

Overall, we have found that for each of the major cargo transport motors, kinesin-1, kinesin-3, and dynein, there is at least one MAP that inhibits each motor and at least one MAP that allows for each motor to progress unimpeded along the microtubule lattice (Figure 5D). These results provide one explanation for why the neuron requires MAPs be compartmentally organized, but also exhibit overlapping spatial and temporal patterns. In addition, the effects of MAPs on motor landing and motility we report here are dramatic, suggesting a strong potential of the MAP code in the direction of motor transport in vivo. We speculate that tubulin modifications may further dictate MAP binding, and thus indirectly gate motor access to the microtubule. This will be a fascinating direction for future work.

It is an outstanding question how motors are spatially distributed into specific neuronal compartments. Three MAPs that localize to the dendrites all inhibit kinesin-1, but allow for kinesin-3 motility on the microtubule, suggesting neurons employ multiple modalities to specifically allow kinesin-3, but prevent kinesin-1 transport into dendrites. In vivo, loss of DCX or DCLK1 impedes the ability of kinesin-3 to transport cargo within dendrites ^6,46^. Our results suggest this effect is not due to direct recruitment or facilitation of kinesin-3 by these MAPs. For example, removal of DCX or DCLK1 could affect the distribution of other MAPs, such as MAP2 and tau that both inhibit kinesin-3 motility. Or these MAPs could affect the lattice itself or the accumulation of posttranslational modifications that could have a cascade of effects on motor transport. The distinct distal localization patterns of DCX and DCLK1 suggest that they are unlikely candidates for the spatial regulation of kinesin-1 and kinesin-3 movement. Our results, combined with other studies, indicate that MAP9 and non-canonical MAPs, such as septins ^56^, may be more important for facilitating kinesin-3 transport into dendrites and excluding kinesin-1.

Our data are consistent with the notion that MAP9 enables kinesin-3 motility on the lattice via a transient ionic interaction with its K-loop. Co-binding analyses between MAPs lead us to hypothesize that MAP9 binds within the interprotofilament valley, but based on its inhibition of dynein, it may also make contacts on the protofilament ridge, overlapping with the kinesin and dynein motor domain binding footprints. Alternatively, the MAP9 projection domain may lead to steric hindrance of motors, irrespective of its binding footprint on the microtubule lattice. High-resolution structural analysis of MAP9 on microtubules will be needed to fully answer this question.

It is tempting to speculate that MAP binding could designate single microtubules, or even sets of protofilaments, as specific tracks for anterograde or retrograde transport. This type of organization, analogous to emerging results in intraflagellar transport ^59^, would prevent collisions between motors and their cargoes and could conceivably allow for MAPs or other factors to independently modulate transport in either direction. Further, while MAP9 can facilitate kinesin-3 and MAP7 can recruit kinesin-1, our data show that under saturating conditions with a simultaneously bound iMAP, these motors are still inhibited. Thus, we speculate that subsets of microtubules or protofilaments must be devoid of iMAPs to facilitate motor transport. Certain MAP properties, such as the cooperative binding exhibited by tau and DCX, could be necessary to establish and maintain homotypic MAP zones on the lattice. Such effects could be further enhanced by potential influences of MAPs on the underlying architecture of the microtubule lattice. Our results therefore provide a basis for understanding how cells can utilize MAPs to establish polarized transport, and ensure an efficient bidirectional transport system.

Finally, this study has implications beyond polarized neuronal transport to any cell or process that relies on MAPs and molecular motors. While some of the MAPs in this study are specifically expressed in the nervous system, many of these MAPs are found in other cell types, such as muscles (tau, MAP7), or in specialized cellular processes, such as mitosis (MAP7, MAP9, DCLK1). It is therefore important to consider how these MAPs affect different types of motors when analyzing any system in which they must share a common microtubule lattice. Further work will be necessary to elucidate how MAPs co-decorate the microtubule in vivo to dictate access by specific motors.

## ACKNOWLEDGEMENTS

This work was supported by the March of Dimes Basil O’Connor Award, NIH grant 1R00HD080981, Pew Scholar Research Award, and Simons Foundation SFARI Award to K.M.O.M, and March of Dimes Basil O’Connor Award and NIH grants 5R00NS089428 and 1R35GM124889 to R.J.M. This material is based upon work supported by the National Science Foundation Graduate Research Fellowship Program under Grant No. 1650042 to B.Y.M. Any opinions, findings, and conclusions or recommendations expressed in this material are those of the authors and do not necessarily reflect the views of the National Science Foundation.

## AUTHOR CONTRIBUTIONS

B.Y.M. and K.M.O.M. conceived of the project and designed the experiments. B.Y.M., J.M.O., and T.T. purified the recombinant proteins. B.Y.M. performed the in vitro TIRF-M experiments and the pull-down assays. R.J.M. and B.Y.M. performed the in vitro TIRF-M experiments with K420 and KIF1A with DCX. R.J.M. and K.M.O.M. performed the in vitro TIRF-M experiments with DDB. B.Y.M. and K.M.O.M. analyzed the in vitro data. B.Y.M. and K.M.O.M. performed the neuronal culture experiments. J.H. and S.S. isolated and cultured the mouse neurons. D.W.N. created the molecular models.

## METHODS

### Molecular Biology

The cDNAs for protein expression in this study were as follows: human Tau-2N4R (Addgene #16316), human MAP7 (GE Dharmacon MGC Collection #BC025777), human MAP2 (Transomics #BC172263), human MAP9 (Transomics # BC146864), human DCX (Addgene #83928), mouse DCLK1 (Transomics #BC133685), human KIF5B (aa 1-560; a gift from R. Vale), and human KIF1A (aa 1-393; Addgene # 61665). Tau-2N4R, MAP2 MAP7, MAP9, and DCLK proteins were cloned in frame using Gibson cloning into a pET28 vector with an N-terminal strepII-Tag, mTagBFP or a superfolder GFP (sfGFP) cassette. DCX proteins were cloned in frame using Gibson cloning into a pET28 vector with a C-terminal sfGFP cassette. K560, K420, and KIF1A, were cloned in frame using Gibson cloning into pET28 vector with a C-terminal mScarlet-strepII cassette. KIF1A _KK>AA_, KIF1A_K-loop>A-loop_, KIF5B_K-insert_, and MAP9_EEE>KKK_ mutations were introduced by overlapping PCR and Gibson cloning into pET28 vector with a C-terminal mScarlet-strepII cassette.

### Protein Expression and Purification

Tubulin was isolated from porcine brain using the high-molarity PIPES procedure as previously described ^60^. For bacterial expression of sfGFP-tau, sfGFP-MAP2, sfGFP-MAP7, mTagBFP-MAP7, sfGFP-MAP9, mTag-BFP-MAP9, sfGFP-MAP9_EEE>KKK_, DCX-sfGFP, sfGFP-DCLK, K560-mScarlet, K420-mScarlet, and KIF1A-mScarlet, KIF1A _KK>AA_-mscarlet, KIF1A_K-loop>A-loop_-mscarlet, KIF5B_K-insert_-mscarlet, BL21-RIPL cells were grown at 37°C until ∼O.D. 0.6 and protein expression was induced with 0.1 mM IPTG. Cells were grown overnight at 18°C, harvested, and frozen. Cell pellets were resuspended in lysis buffer (50 mM Tris pH 8, 150 mM K-acetate, 2 mM Mg-acetate, 1 mM EGTA, 10% glycerol) with protease inhibitor cocktail (Roche), 1 mM DTT, 1 mM PMSF, and DNAseI. Cells were then passed through an Emulsiflex press and cleared by centrifugation at 23,000 x *g* for 20 mins. Clarified lysate from bacterial expression was passed over a column with Streptactin Superflow resin or Streptactin XT Superflow resin (Qiagen). After incubation, the column was washed with four column volumes of lysis buffer, then bound proteins were eluted with 3 mM desthiobiotin (Sigma) or 50 mM D-biotin (CHEM-IMPEX) in lysis buffer. Eluted proteins were concentrated on Amicon concentrators and passed through a superose-6 (GE Healthcare) gel-filtration column in lysis buffer using a Bio-Rad NGC system. Peak fractions were collected, concentrated, and flash frozen in LN_2_. Protein concentration was determined by measuring the absorbance of the fluorescent protein tag and calculated using the molar extinction coefficient of the tag. The resulting preparations were analyzed by SDS polyacrylamide gel electrophoresis (SDS-PAGE).

### Pull-down Assays

Pull-down assays were performed with either sfGFP-DCLK or sfGFP-MAP9 tagged at the C-terminal end with a FLAG epitope. FLAG beads (Thermofisher) were washed into assay buffer (100 mM Tris, pH 7.4, 150 mM NaCl, 4 mM MgCl, 1 mM EGTA, 0.5% TX-100, supplemented with 1 mM DTT and 4 mM ATP), then incubated with 500 nM DCLK, 500 nM MAP9, or buffer (beads alone control) for 1 hour rotating at 4°C. FLAG beads were then washed in assay buffer five times, then resuspended in assay buffer and 500 nM for KIF1A was added to the beads alone control and the experimental conditions. The 350 μL final volume solutions were incubated for 1 hour rotating at 4°C. The supernatants were collected, then the bead pellets were washed five times in assay buffer and resuspended in one bead bed volume. Gel samples of the supernatants and pellets were analyzed by SDS-PAGE. The supernatant samples that were run on SDS-PAGE were 15% of the pellet samples.

### TIRF Microscopy

For TIRF-M experiments, a mixture of native tubulin, biotin-tubulin, and fluorescent-tubulin purified from porcine brain (∼10:1:1 ratio) was assembled in BRB80 buffer (80mM PIPES, 1mM MgCl_2_, 1mM EGTA, pH 6.8 with KOH) with 1mM GTP for 15 min at 37°C, then polymerized MTs were stabilized with 20 μM taxol. Microtubules were pelleted over a 25% sucrose cushion in BRB80 buffer to remove unpolymerized tubulin. Flow chambers containing immobilized microtubules were assembled as described^61^. Imaging was performed on a Nikon Eclipse TE200-E microscope equipped with an Andor iXon EM CCD camera, a 100X, 1.49 NA objective, four laser lines (405, 491, 568, and 647 nm) and Micro-Manager software ^62^. All experiments were performed in assay buffer (60 mM Hepes pH 7.4, 50 mM K-acetate, 2 mM Mg-acetate, 1 mM EGTA, and 10% glycerol) supplemented with 0.1mg/mL biotin-BSA, 0.5% Pluronic F-168, and 0.2 mg/mL κ–casein (Sigma) and 10 uM taxol.

For all MAP plus motor experiments, saturating concentrations for the MAPs and single molecule concentrations for the motors were premixed and flowed into the chamber at the same time. For the MAP competition plus motor experiments, saturating concentrations of the MAPs and single molecule concentrations for the motors were premixed and flowed into the chamber at the same time. For live imaging, images were taken every 0.5 seconds for a total of 4 minutes for K560 (K420) and every 0.24 seconds for a total of 2 minutes for KIF1A (KIF1A _KK>AA_, KIF1A_K-loop>A-loop_, KIF5B_K-insert_). Kymographs were made from movies of K560, K420, KIF1A, KIF1A _KK>AA_, KIF1A_K-loop>A-loop_, and KIF5B_K-insert_ in the absence or presence of MAPs and landing rates and velocity parameters were measured for individual events and runs, respectively. Velocity data were fit with a Gaussian equation.

For all competition experiments, proteins were premixed at equimolar concentrations and flowed into the chamber at the same time. Competition experiments were performed with a mixture of either mTagBFP-MAP9 and sfGFP-DCX or sfGFP-DCLK or mTagBFP-MAP9 and sfGFP-tau or sfGFP-MAP7.

For all saturation curves, a concentration series was performed for each protein. For fluorescence intensity analysis, ImageJ was used to draw a line across the microtubule of the MAP channel and the integrated density was measured. The line was then moved adjacent to the microtubule of interest and the local background was recorded. The background value was then subtracted from the value of interest to give a corrected intensity measurement. The fluorescence intensity data were fit with a one site binding hyperbola equation to derive the K_D_ for each MAP.

### Immunocytochemistry

For neuronal cultures, hippocampi were dissected out from E16.5-18.5 mouse embryonic brains and hippocampal neurons were isolated using Worthington Papain Dissociation System (Worthington Biochemical Corporation) following manufacturer’s protocol. Primary neurons were plated at density of 100k neurons per glass coverslips coated with 0.05 mg/ml Poly-D-Lysine (mol wt 70,000-150,000, Millipore-Sigma) and cultured in Neurobasal media supplemented with glucose, GlutaMAX (ThermoFisher), B27 and P/S. Media was changed every two days until neurons were fixed at DIV4. The cultures were fixed in 4% paraformaldehyde for 20 minutes at room temperature, washed several times with PBS, permeabilized in PBS with 0.3% Triton X-100 (PBS-TX), and blocked with 5% BSA in PBS for 1 hour at room temperature. Cultures were then incubated overnight at 4°C with primary antibodies at a concentration of 1:500 for rabbit anti-MAP7 (Thermofisher PA5-31782), 1:500 for mouse monoclonal anti-Tau-1 (Millipore MAB3420), 1:500 anti-chicken tau (Genetex GTX49353), 1:500 anti-rabbit MAP2 (Sigma M3696), 1:500 anti-rabbit MAP9 (Invitrogen PA5-58145), 1:500 anti-mouse DCLK (Invitrogen MA5-26800), 1:500 anti-mouse DCX (Invitrogen MA5-17066), 1:1000 for mouse anti-alpha Tubulin (Sigma Clone DM1A T9026), or 1:1000 for rabbit anti-beta Tubulin (Abcam ab6046). Secondary antibodies were used at 1:1000 for Cy3 donkey anti-rabbit, Cy5 donkey anti-mouse, or Cy3 goat anti-chicken and incubated for 1 hour at room temperature. Cells were then rinsed several times with PBS and mounted using VectaShield mounting medium (Vector Laboratories). Coverslips were imaged on a Leica SPE laser scanning confocal microscope using an oil immersion 60x or 100x objective.

### Statistical Analysis

All statistical tests were performed with a two-tailed unpaired Student’s t-test.

### Data Availability

The data that support the findings of this study are available from the corresponding author upon reasonable request.

## Supplementary Figures and Figure Legends

**Figure S1.**
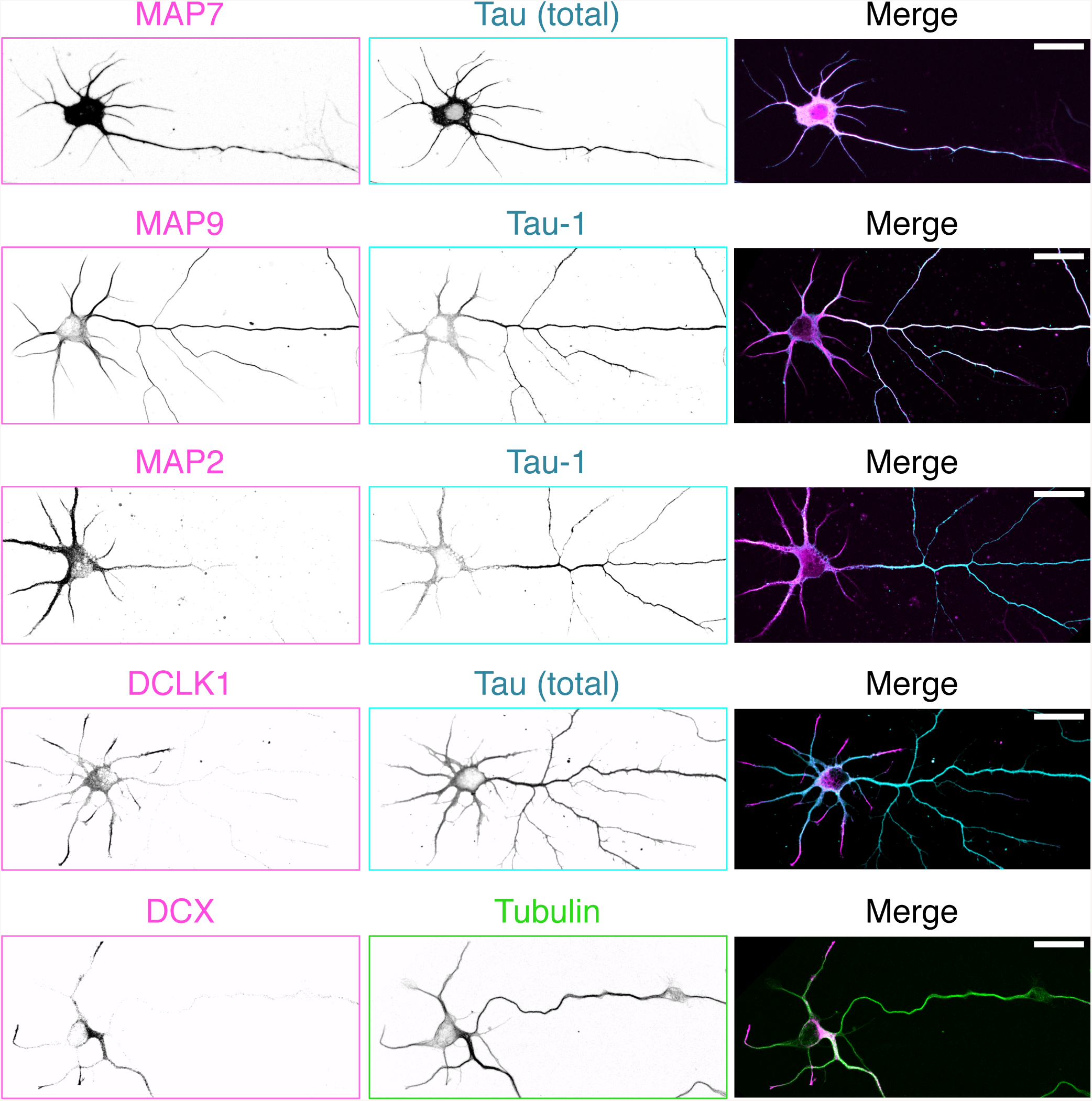
Localization patterns of MAPs in DIV4 mammalian neurons. Immunocytochemistry of mouse DIV4 neuronal cultures with antibodies against dephospho-tau (Tau-1), total tau, MAP7, MAP2, MAP9, DCLK1, DCX, and tubulin. Scale bars are 25 µm. n = three independent neuronal cultures for all conditions.

**Figure S2.**
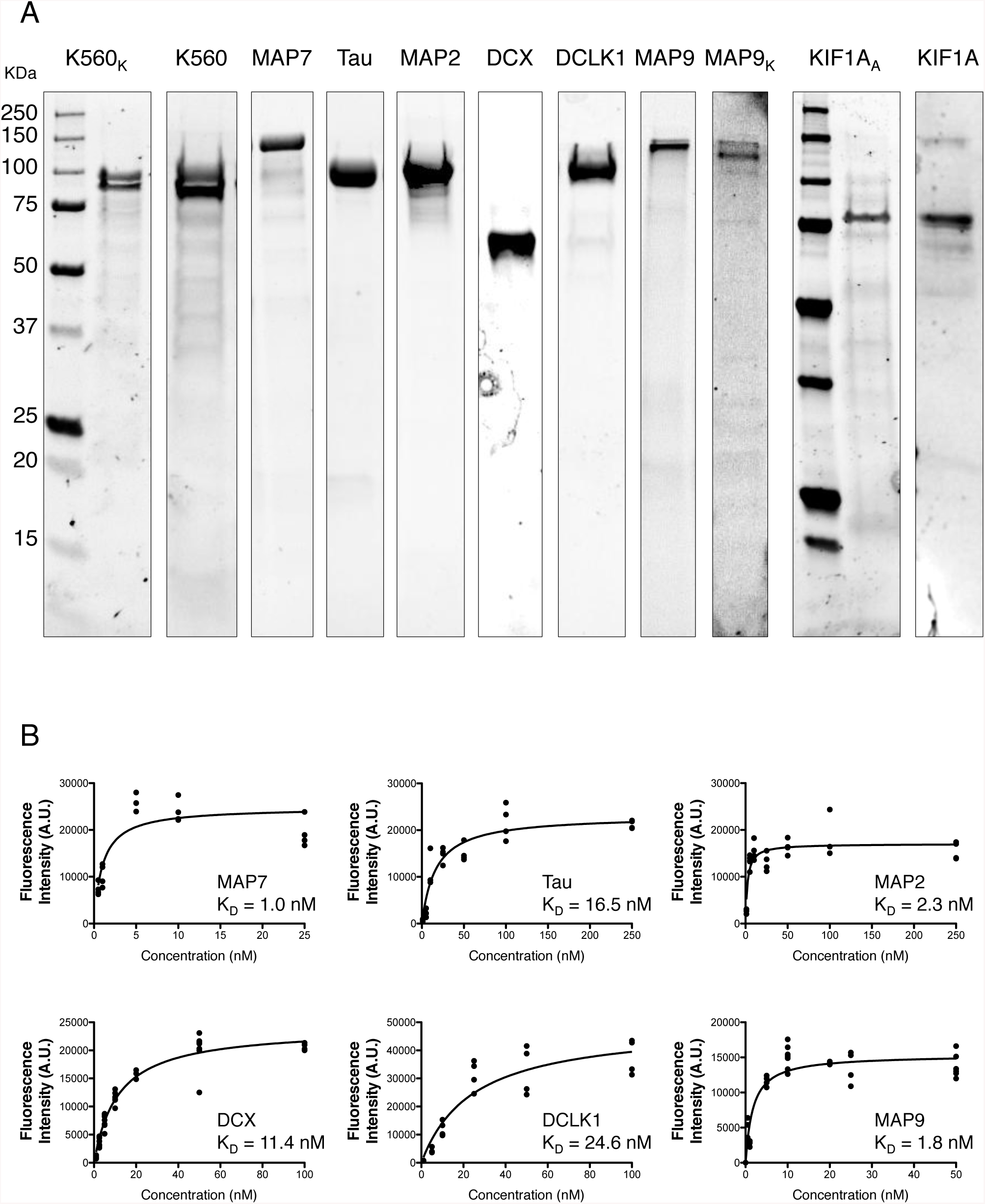
Gels and saturation curves for purified recombinant proteins used in this study. **(A)** Coomassie Blue-stained SDS-PAGE gels of K560K-mScarlet, K560-mScarlet, sfGFP-MAP7, sfGFP-tau, sfGFP-MAP2, DCX-sfGFP, sfGFP-DCLK1, sfGFP-MAP9, sfGFP-MAP9_EEE>KKK_, KIF1A_KK>AA_-mScarlet, and KIF1A-mScarlet. **(B)** Quantification of fluorescence intensity of microtubule-bound MAPs (all tagged with sfGFP) plotted against concentration. The K_D_ for each MAP are indicated.

**Figure S3.**
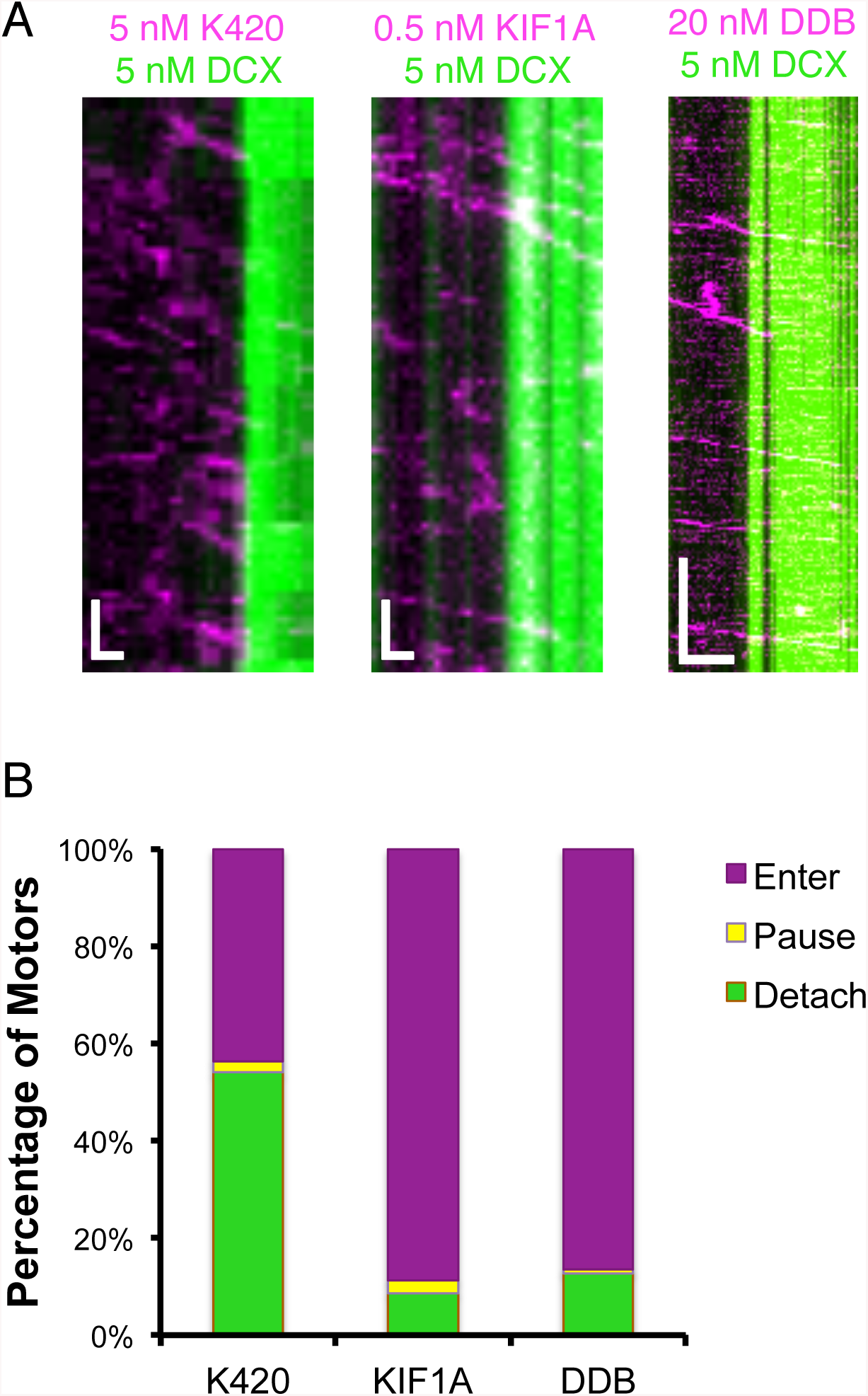
DCX patch assemblies differentially affect kinesin-1, kinesin-3, and dynein motors traveling along a microtubule. **(A)** Kymographs depicting K420-mScarlet (kinesin-1), KIF1A-mScarlet (kinesin-3), and DDB-TMR (dynein-dynactin-BicD) motors encountering a DCX-sfGFP patch. **(B)** Quantification of the percent of K420, KIF1A and DDB motors that detach, pause, or pass DCX patches. For K420 motors, 54.0 % detach, 2.3 % pause, and 43.7 % enter DCX patches (174 events from two independent experiments). For KIF1A motors, 8.7 % detach, 2.6 % pause, and 88.7 % enter DCX patches (115 events from two independent experiments). For DDB motor complexes, 12.8 % detach, 0.7 % pause, and 86.5 % enter DCX patches (133 events from two independent experiments).

**Figure S4.**
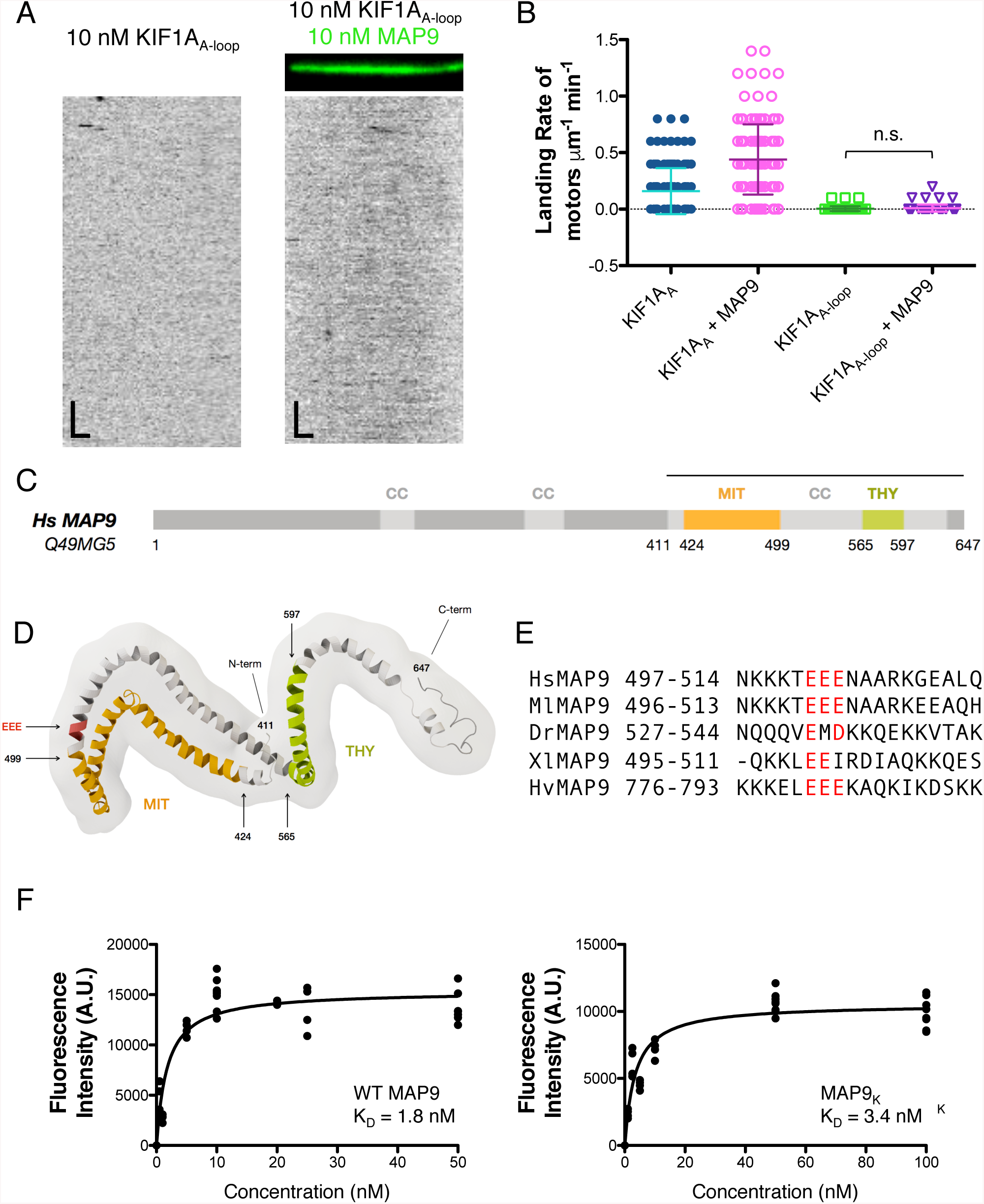
Dissection of the MAP9 interaction with kinesin-3. **(A)** TIRF-M images and kymographs of 10 nM KIF1A_A-loop_-mScarlet + 1 mM ATP in the absence and presence of 10 nM sfGFP-MAP9 (green). Images are 10.3 μm wide. Scale bars: 1 μm (*x*) and 5 sec (*y).* **(B)** Quantification of the landing rates of 5nM KIF1A_A-loop_-mScarlet compared with KIF1A_A_-mScarlet + 1 mM ATP in the absence and presence of MAP9. KIF1A_A_ data are reproduced from Figure 3 for comparison. Means ± s.d. for motors μm^-1^ min^-1^ are: 0.005 ± 0.02 for KIF1A_A-loop_ alone (n = 103 kymographs from two independent trials) and 0.009 ± 0.03 for KIF1A_A-loop_ + MAP9 (n = 103 kymographs from two independent trials). All datapoints are plotted with lines indicating means ± s.d. *P* = 0.26 using a student’s t-test for KIF1A_A-loop_ *vs.* KIF1A_A-loop_ + MAP9. **(C)** Domains and motifs of human MAP9. CC: coiled-coil regions. MIT: microtubule interacting and trafficking domain. THY: thymosin beta actin-binding motif (Venoux et al., 2008). **(D)** Model of the MAP9 C-terminus (aa 411-647), predicted by *de novo* and homology modeling (Buchan et al., 2010). **(E)** Sequence alignment between HsMAP9, *Myotis lucifugus* MAP9 (MlMAP9), *Danio rerio* MAP9 (DrMAP9), *Xenopus laevis* (XlMAP9), and *Hydra vulgaris* MAP9 (HvMAP9). It is important to note that hydra MAP9 is ∼300 aa longer at the amino-terminus than human MAP9, thus this region corresponds to the same region of the microtubule-binding domain of human MAP9. **(F)** Quantification of fluorescence intensity of microtubule-bound sfGFP-MAP9 or sfGFP-MAP9_EEE>KKK_ plotted against concentration. The K_D_ for the wild type and mutant MAP9 proteins are indicated.

**Figure S5.**
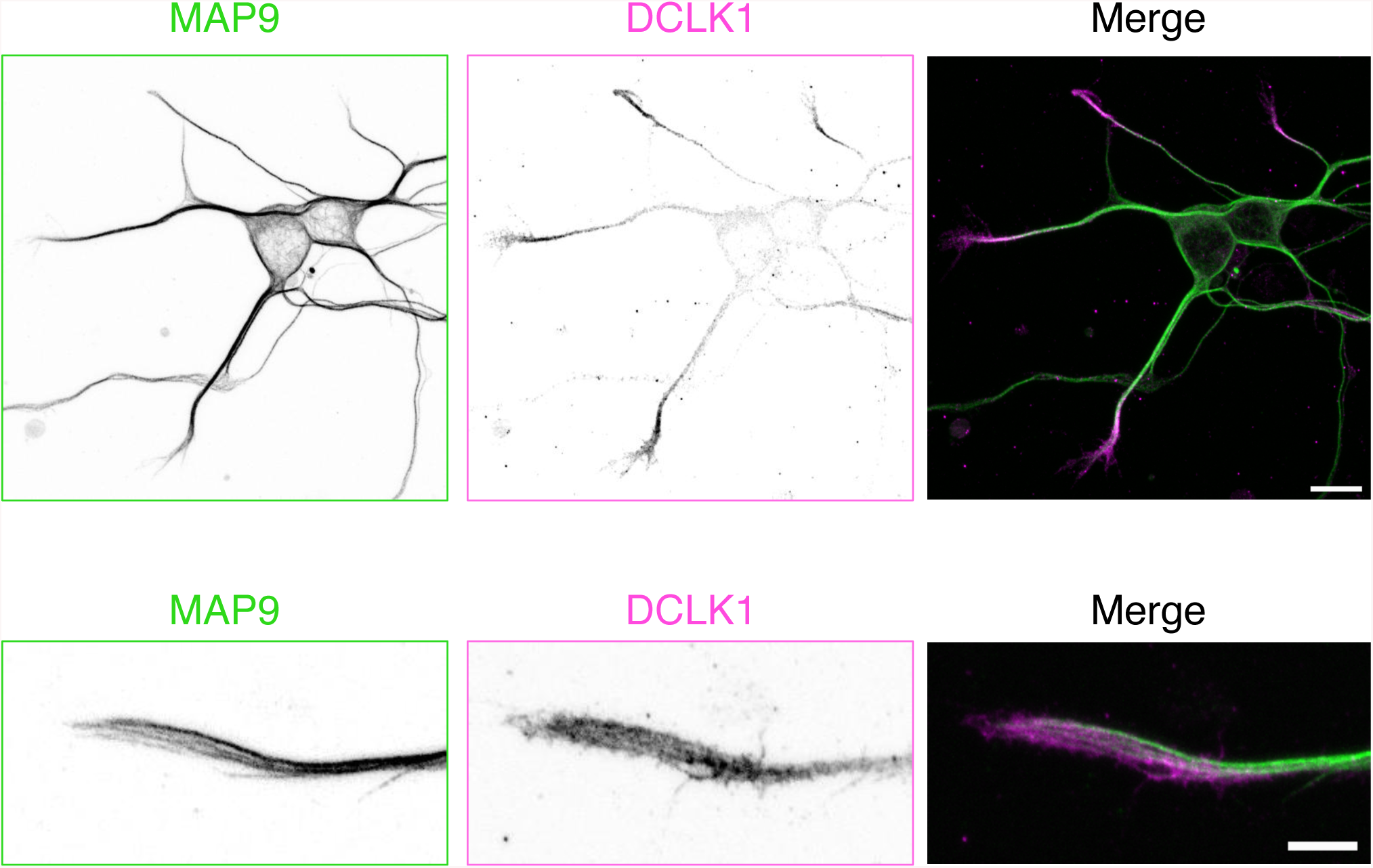
MAP9 and DCLK1 are inversely correlated in distal dendrites. Immunocytochemistry of mouse DIV4 neuronal cultures with antibodies against MAP9 and DCLK1. Scale bars are 10 µm for the large image and 5 µm for the zoomed-in image. n = two independent neuronal cultures.

